# Generating Joint Transcriptomic and Morphological Responses to Drug Perturbations via Rectified Flow

**DOI:** 10.64898/2026.02.02.703189

**Authors:** Shourya Verma, Mengbo Wang, Luopin Wang, Simran Kadadi, Mario Sola, Majid Kazemian, Ananth Grama, Nadia Atallah Lanman

**Affiliations:** Department of Computer Science, Purdue University; Purdue Institute for Cancer Research, Purdue University; Department of Comparative Pathobiology, Purdue University

**Keywords:** Perturbation Modeling, Generative AI, Multi-modal Learning, Transcriptomics, Cell Painting

## Abstract

**Motivation:** Predicting cellular responses to drug perturbations requires capturing complex dependencies between transcriptomic and morphological changes. Existing approaches model these modalities in isolation, missing critical molecular-phenotypic relationships that occur simultaneously during drug treatment. No current method jointly predicts gene expression profiles and cellular morphology from chemical perturbations.

**Results:** We introduce PertFlow, a unified computational framework that simultaneously predicts treatment gene expression (bulk RNA-seq) and generates cellular morphology (Cell Painting images) from control cellular states, conditioned on drug metadata. Evaluated on paired RNA-seq and imaging data from 3 cell lines and 40 compounds (17,242 samples), PertFlow achieves Pearson correlation of 0.780±0.264 for transcriptomic prediction and FID of 24.06 for morphological generation. Board-certified pathologists rated generated images with median similarity scores of 7.11-7.89/10. The model successfully recovers known drug mechanisms including microtubule disruption, DNA damage, and MAPK pathway inhibition, with gene enrichment analysis confirming activation of expected biological pathways (EMT, p53, apoptosis).

**Availability:** Code and pretrained models will be available at https://github.com/wangmengbo/PertFlow.

## Introduction

Understanding how drugs alter cellular states is essential for drug discovery, mechanistic understanding, and personalized medicine. High-throughput profiling now enables paired RNA sequencing and imaging, offering complementary insights: transcriptomics captures molecular mechanisms and gene regulation, while morphology reflects structural and phenotypic changes. These modalities are intrinsically linked, as gene expression drives morphological transformations, and structural changes modulate gene activity, yet most models treat them in isolation.

Existing transcriptomics-focused methods predict gene expression responses to perturbations but cannot model morphological effects. Foundational approaches like scGen (1) pioneered VAE-based latent space arithmetic for unseen perturbation combinations, with CPA (2) and chemCPA (3) extending compositional modeling to chemical structures for completely unseen drugs. GEARS (4) applies geometric deep learning on gene interaction graphs for genetic perturbations, while PRnet (5) and TranSiGen (6) employ generative models for novel compounds, though the latter is limited to denoising and reconstruction. Recent evaluations (7) suggest deep learning models do not significantly outperform traditional baselines for genetic perturbations, leaving chemical perturbation effects largely unexplored.

In parallel, imaging and cross-modal integration approaches emphasize prediction or measurement over generation. Cell Painting (8) enables multiplexed morphological profiling, with tools like CellProfiler (9) and Cellpose (10) advancing analysis and segmentation through deep learning (11). Recent generative efforts focus on image-to-image translation: PhenDiff (12) uses conditional diffusion models and (13) employs conditional GANs, but both operate without transcriptomic context. Cross-modal frameworks like BLEEP (14), SCHAF (15), and TransformerST (16) predict spatial gene expression from histology, while Perturb-multi-modal (17), CRISPR ST (18), and fusion-based methods (19) integrate imaging and RNA-seq for genetic perturbations but focus on measurement rather than synthesis. Critically, no method jointly generates both transcriptomic and morphological responses to chemical perturbations.

Joint multi-modal generation poses three fundamental challenges: (1) aligning transcriptomic and morphological data across fundamentally different representational spaces; (2) capturing complex drug conditioning involving compound structure, dose, cell type, and timepoint; and (3) simultaneously predicting discrete gene expression and continuous image data with biological realism and cross-modal consistency. While generative frameworks show promise, diffusion models achieve state-of-the-art performance in image, protein, and molecule generation (20, 21), and rectified flow (22, 23) offers computational efficiency through straight-line transport between distributions, they have not been applied to joint cellular response prediction from chemical perturbations.

We introduce **PertFlow** (Figure 1), a unified framework that jointly predicts treatment gene expression and synthesizes cellular morphology from control conditions, conditioned on drug metadata. PertFlow integrates control RNA-seq, control images, and drug information through cross-modal attention into a shared embedding space, then employs regression for RNA-seq prediction and generative modeling for morphology synthesis with cross-modal consistency constraints. Our approach enables accurate predictions from complete multi-modal inputs or single-modality data alone. Evaluated on paired bulk RNA-seq and Cell Painting imaging from 3 cell lines and 40 compounds (17,242 samples), PertFlow achieves strong performance (Pearson r=0.780±0.264, FID=24.06), recovers known drug mechanisms (microtubule disruption, DNA damage, MAPK inhibition), and demonstrates biological accuracy through pathologist evaluation (7.11-7.89/10) and gene enrichment analysis. To our knowledge, PertFlow is the first method capable of joint transcriptomic and morphological response prediction for chemical perturbations, which could enable downstream applications in virtual drug screening, mechanism discovery, and integrated pharmacological modeling.

**Fig. 1.**
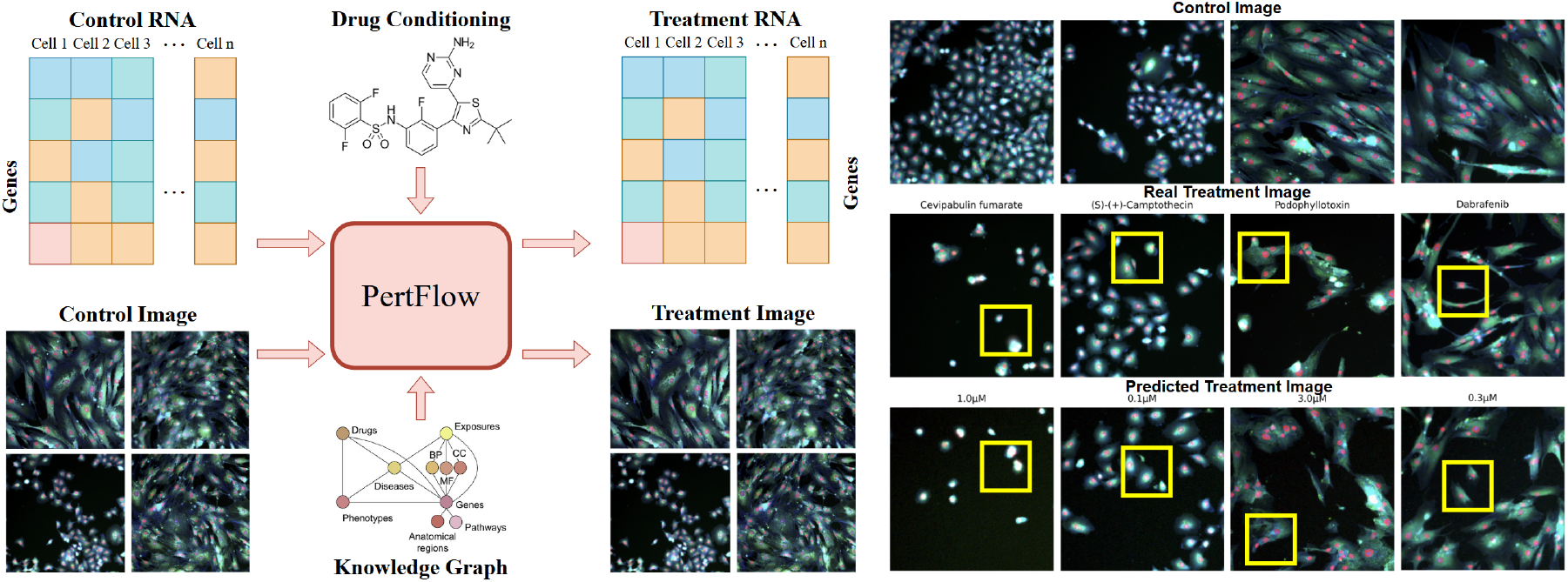
(LEFT) Cross-modal mapping from control RNA-seq and image to treatment RNA-seq and image with drug conditioning through PertFlow. (RIGHT) Comparison of generated treatment vs real treatment images with drug name and concentration. Yellow boxes indicate similar features.

## Methods

### Problem Formulation

We formalize the drug conditioned multi modal generation problem as learning a mapping from control cellular states to treatment responses across transcriptomic and morphological modalities. Given control gene expression 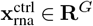 where *G* is the number of genes, control cellular images 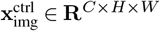 with *C* channels and spatial dimensions *H* ×*W*, and drug conditioning information **c** = {*c*_compound_, *c*_cell_, *c*_conc_, *c*_time}_ including compound identity, cell line, concentration, and timepoint, our objective is to generate treatment outcomes, where *f*_*θ*_ represents our unified generative model parameterized by *θ*:

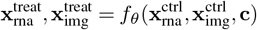

### Dataset Description

Our study leverages the Ginkgo Data Platform (GDP) series (24), a multimodal dataset integrating transcriptomic profiles (GDPx1/GDPx2) and four-channel fluorescence microscopy images (GDPx3) from drug-treated cell cultures. We curated paired bulk RNA-seq and Cell Painting imaging dataset from 3 cell lines and 40 drugs as illutrated in Figure 2 (RIGHT). We implemented cross-modal pairing by identifying overlapping compounds and experimental conditions, standardizing metadata (concentration units, cell line nomenclature, temporal alignment), and establishing DMSO controls as baseline references. Transcriptomic preprocessing follows established protocols like total count normalization to 10^6^ reads per sample, log1p transformation, and highly variable gene selection (n=8000) using scanpy, to focus on the most informative genomic features. Image preprocessing addresses 16-bit microscopy data through proper intensity scaling (16-bit to [-1,1] range), percentile-based contrast enhancement (1^*st*^-99^*th*^ percentile) applied per channel, and bilinear interpolation to uniform spatial dimensions. The dataset consists of 17242 paired samples that were split by 80:20 for training and testing.

**Fig. 2.**
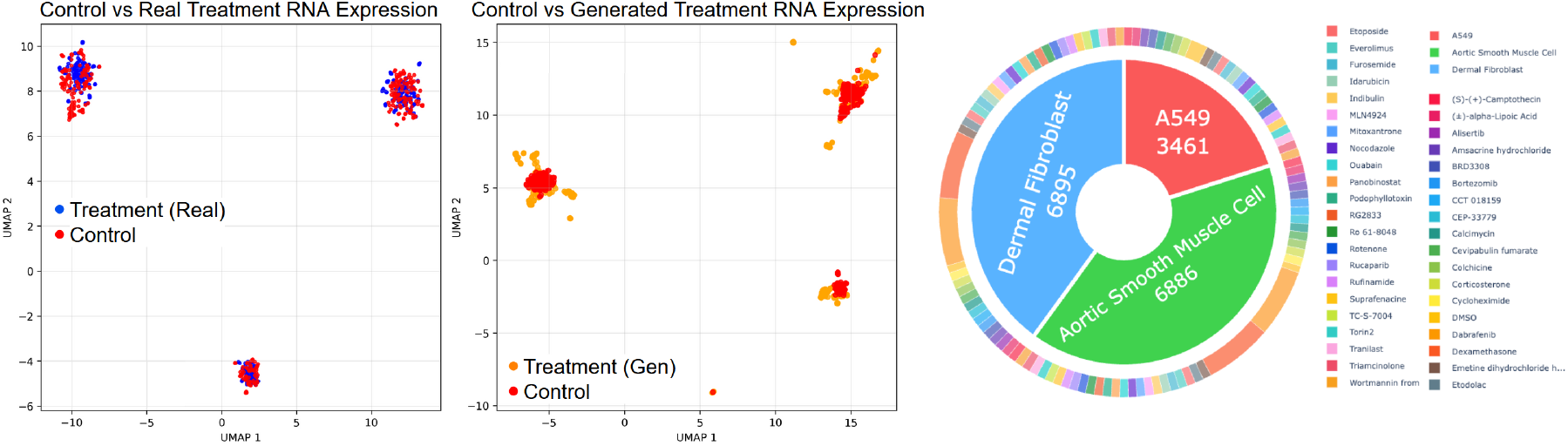
(LEFT) UMAP representation of control vs real treatment vs generated treatment gene expression data. (RIGHT) Distribution of cell lines and compounds in dataset.

Drug compounds are represented through multi-modal molecular encodings combining structural and physicochemical information. We extract Morgan and RDKit molecular fingerprints (1024 bits each) from canonical SMILES strings, providing binary structural descriptors capturing sub-structural patterns and pharmacophoric features. Molecular descriptors include eighteen 2D properties (molecular weight, logP, topological polar surface area, hydrogen bond donors/acceptors, rotatable bonds, aromatic rings, and complexity measures) and five 3D properties when available from SDF structures. For compounds lacking preprocessed molecular data, we implement on-demand SMILES processing with RDKit to ensure comprehensive coverage. Missing molecular information is handled through zero-padding with appropriate masking, while molecular descriptors are normalized using dataset-wide statistics to ensure stable training dynamics across diverse chemical spaces. The dataset allows stratified paired control-treatment comparisons, where drug-treated samples are systematically matched with vehicle DMSO controls from identical cell lines and experimental conditions. This preserves the combinatorial structure of cell line-compound-dose-time relationships across training and validation partitions, ensuring robust model generalization across the full experimental parameter space.

### Architecture

PertFlow (Figure 3) uses a shared representation learning paradigm with 3 components: (1) individual modality encoders that process control RNA-seq and imaging data, (2) cross-modal attention mechanism that aligns features across modalities, (3) generation heads that produce treatment RNA-seq via direct prediction and treatment images via rectified flow dynamics.

**Fig. 3.**
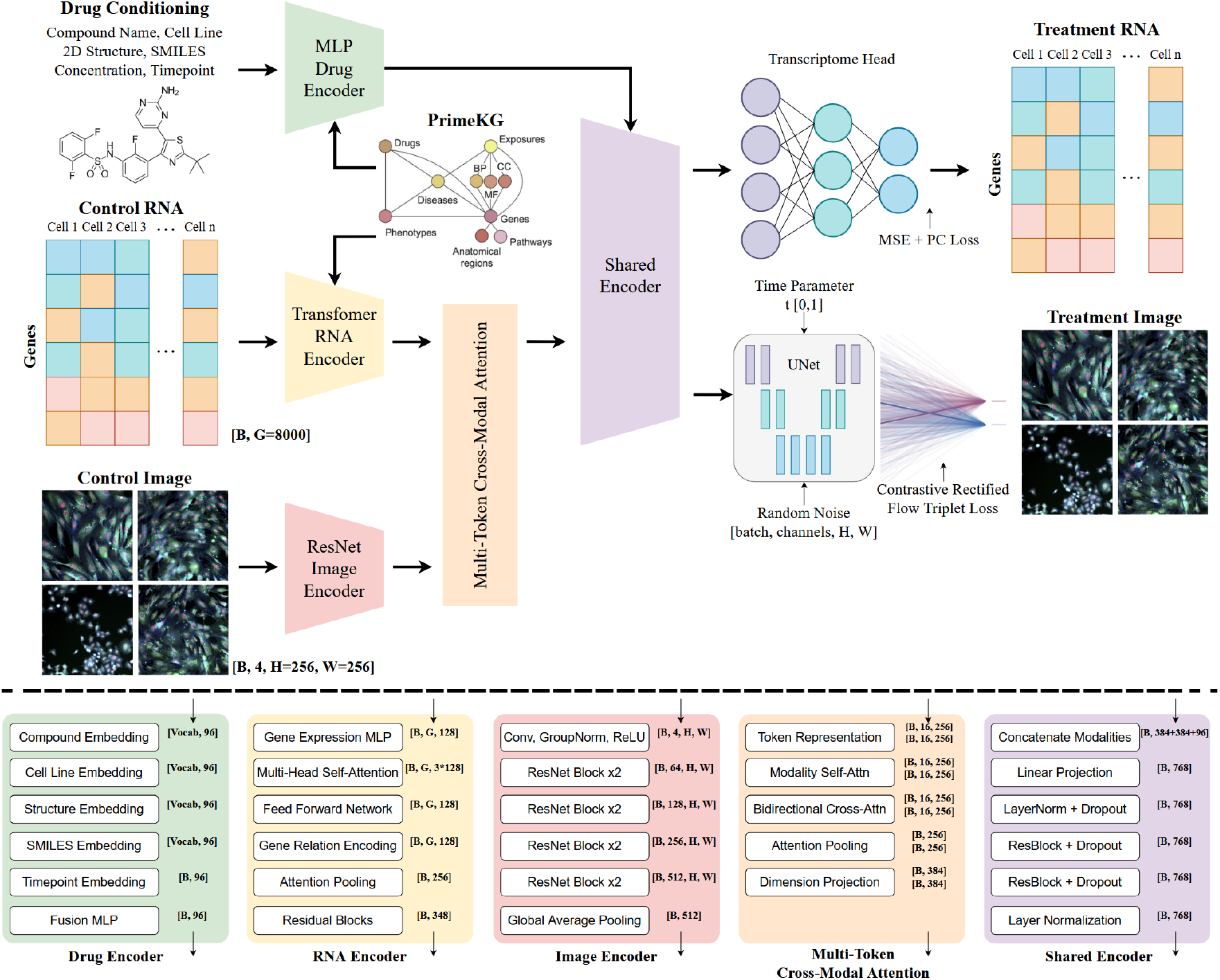
PertFlow architecture for drug conditioned control RNA-image to treatment RNA-image. Input RNA-seq and image going through their respective encoders; output from the two encoders then pass through the multi-token cross-modal attention, before entering the shared encoder along with conditioning information which passes through the drug encoder. The transcriptome head uses MSE loss to predict treatment RNA, and the image UNet (25) with noise and time parameter input uses triplet contrastive loss and rectified flow dynamics to generate treatment cellular image, from the shared embeddings, respectively.

PertFlow processes control RNA-seq and cellular images through specialized encoders. The RNA-seq encoder applies multi-layer self-attention to capture gene-gene interactions, embedding each gene expression value into high-dimensional space and processing through attention and feedforward layers with residual connections. Attention-weighted pooling aggregates the final embeddings into a single RNA-seq representation encoding the transcriptomic state. Control images are encoded through a ResNet-style convolutional architecture with global pooling, extracting hierarchical visual features from cellular morphology. Both modalities are enhanced through PrimeKG knowledge graph integration. For RNA-seq, a heterogeneous graph neural network processes protein-protein interactions and pathway information. For drugs, the graph encoder processes drug-protein interactions and pharmacological relationships. Knowledge embeddings are integrated additively with learned representations using weighting factors *α*_drug_ = 0.3 and *α*_RNA_ = 0.3. Drug conditions are encoded through a fusion module combining learned embeddings for compound and cell line identities with encoded concentration and time parameters.

Each modality embedding is projected to *K* token representations to prevent information bottlenecks. RNA and image tokens undergo self-attention within modalities, then bidirectional cross-attention: RNA tokens attend to image tokens and vice versa, integrating information across modalities. Cross-attended tokens combine with original tokens through residual connections, then aggregate to single vectors via attention pooling. The enhanced RNA and image embeddings concatenate with drug conditioning and process through a shared encoder, a multi-layer perceptron producing unified representation **h**_shared_ where all modalities converge. This representation branches to two task-specific generation heads. Treatment transcriptomes are generated through direct regression from **h**_shared_ via a fully-connected prediction head. For images, PertFlow adapts rectified flow, defining linear interpolation paths 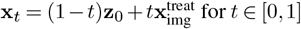 with constant velocity 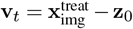. A UNet predicts this velocity field conditioned on noisy state **x**_*t*_, timestep *t*, and **h**_shared_ injected through cross-attention layers. During inference, an adaptive DOPRI5 ODE solver integrates the learned velocity field from noise to treatment image with automatic step size adjustment.

Transcriptome prediction combines MSE with Pearson correlation loss:

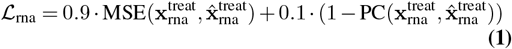

where MSE ensures gene-level accuracy and correlation preserves relative expression patterns. The rectified flow objective trains velocity prediction:

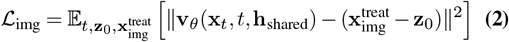

over random timesteps *t* ∼Uniform(0, 1) and noise samples **z**_0_.

Triplet contrastive consistency enforces coherent multi-modal predictions by comparing aligned versus misaligned features:

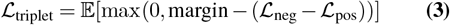

where ℒ_pos_ is prediction error with aligned RNA-image features and ℒ_neg_ with shuffled features. The complete objective combines all losses:

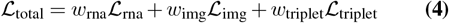

with weights *w*_rna_ = 0.5, *w*_img_ = 0.5, *w*_triplet_ = 0.05. All parameters are jointly optimized for end-to-end multi-modal learning.

### Training Parameters

We use 4 attention heads with embedding dimension 128, applying 1 layer of self-attention followed by attention-based pooling. Multi-token representations use *K* = 16 tokens with hidden dimension 256 and 8 attention heads. The rectified flow UNet uses 192 base channels with channel multipliers (1, 2, 2, 2), attention at 16×16 resolution, and cross-attention conditioning at layers 2, 3, 4, and 5. Models are trained with AdamW optimizer (*β*_1_ = 0.9, *β*_2_ = 0.95), learning rate 10^*−*4^ with cosine annealing, and automatic mixed precision. Cross-modal consistency weights are gradually increased during training to ensure stable convergence. RNA-seq generation requires a single forward pass, while image generation uses 7-10 DOPRI5 steps with relative tolerance 10^*−*3^ and absolute tolerance 10^*−*4^ for high-quality synthesis. The models were trained with an effective batch size of 32 taking 5 hours on 8 H100 NVIDIA GPUs with 80GB VRAM. Further architectural details can be found in the appendix.

## Experiments

We emphasize, since our method introduces the problem of multi-modal RNA-Image generation with drug conditioning, we have **no previous multi-modal method** to fairly compare to as baseline. To create baselines we trained three diffusion models, along with ablations of knowledge graph module, triplet contrastive objective, and Pearson correlation loss, with RNA only and Image only models. We still included PRNet (5) and PhenDiff (12) as uni-modal method for reference. We set the state-of-the-art performance for this problem on the GDPx3 dataset.

### Drug effects on gene expression and cell morphology

We trained PertFlow (control RNA-seq and image to treatment RNA-seq and image), PertRNA (control RNA-seq to treatment RNA-seq), PertImage (control image to treatment image), and their respective ablations omitting the knowledge-graph and contrastive rectified flow objective. PertFlow demonstrates strong performance in predicting treatment gene expression from control conditions in Table 1. Achieving Pearson correlation (0.780 ± 0.264) and Spearman correlation (0.792 ± 0.041) across drug perturbations, measuring with all genes. MSE (0.231 ± 0.708) and MAE (0.110 ± 0.166) indicate robust prediction accuracy for transcriptomic responses. While the PertRNA baseline achieved correlation metrics (Pearson r (0.779 ± 0.081), Spearman r (0.795 ± 0.026)), PertFlow’s joint modeling approach maintains competitive performance while simultaneously generating cellular morphological responses compared to baselines and ablations.

**Table 1.**
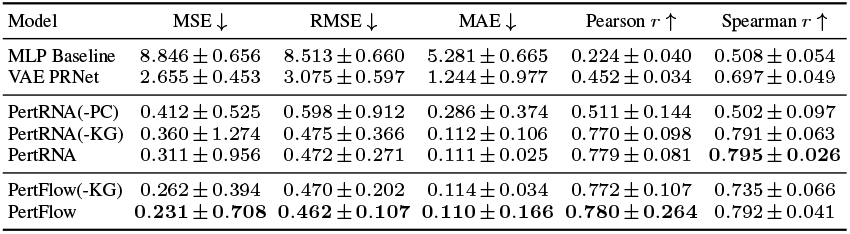
PertFlow & PertRNA metrics (mean ± std)

PRNet (5) is a perturbation-conditioned generative model comprising three components: a Perturb-adapter encod-ing compound SMILES structures to latent embeddings, a Perturb-encoder mapping perturbation effects to latent space, and a Perturb-decoder estimating Gaussian distributions of perturbed transcriptional profiles. The simple MLP baseline replaces the full encoder-decoder architecture with a multi-layer perceptron that directly learns perturbation effects on gene expression using MSE loss. This simplified architecture achieved only Pearson correlation of 0.224±0.040 and Spearman correlation of 0.508±0.054, with MSE of 63.846±0.656. The VAE PRNet baseline implements the complete VAE-inspired framework with encoder-decoder architecture estimating Gaussian distributions parameterized by mean and variance, but operates on transcriptional data alone without multi-modal integration. VAE PRNet substantially improved over the MLP variant with Pearson correlation of 0.452±0.034 and Spearman correlation of 0.697±0.049 (MSE 25.655±0.453), yet remained far below PertRNA (Pearson 0.779±0.081, Spearman 0.795±0.026) and PertFlow (Pearson 0.780±0.264, Spearman 0.792±0.041). Both PRNet variants predict treatment RNA profiles but lack the cross-modal enhancement, knowledge graph integration, and shared representation learning that enables PertFlow to achieve coherent simultaneous prediction of transcriptional and morphological responses while maintaining competitive performance on both modalities.

PertFlow also outperformed the PertImage baseline in the image generation task achieving higher SSIM (0.205 ±0.097), PSNR (11.66 ±0.68), LPIPS (0.511± 0.038), and lower FID (24.06). This demonstrates that incorporating cross-modal information and drug conditioning does not compromise transcriptomic prediction and cellular image generation quality, while enabling the unique capability of joint multi-modal generation. Strong correlation values and lower FID indicates PertFlow successfully captures the complex cellular relationships between drug perturbations and gene expression changes. We compare our PertFlow model with three PertDiff diffusion variants in Table 2. Diffusion serves as the baseline against our contrastive rectified flow method. The standard DDPM formulation trains the model to predict the injected noise *ϵ* ∼*N* (0, *I*) from the noisy input *x*_*t*_. This requires disentangling signal from noise across noise levels, causing instabilities when signal-to-noise ratios are low. The weak performance of PertDiff_*N*_ (SSIM: 0.010, FID: 246.01) illustrates these challenges, as the model fails to generate coherent cellular structures from pure noise predictions. The direct *x*_0_ parameterization predicts the clean target *x*_0_ from noisy *x*_*t*_. While more stable, it forces the model to implicitly learn the full denoising trajectory. PertDiff_*x*0_ achieves better but still limited results (SSIM: 0.192, FID: 73.63), reflecting the lack of strong theoretical grounding. The velocity (*v*) parameterization improves training stability by predicting *v* = *α*_*t*_*ϵ* − *σ*_*t*_*x*_0_, which balances objectives across timesteps, reduces variance, and improves gradient flow. PertDiff_*V*_ shows marked improvement (SSIM: 0.194, FID: 55.92), validating this formulation for biological image generation. Compared to baselines and ablations we observe that PertFlow successfully learns more meaningful representations that generalize across different compounds, concentrations, cell lines, and timepoints in the dataset.

**Table 2.** PertFlow & PertImage metrics (mean ± std)

| Model | SSIM ↑ | PSNR ↑ | LPIPS → | FID ↓ |
| --- | --- | --- | --- | --- |
| UNet Baseline | 0.085 ± 0.013 | 03.55 ± 0.73 | 2.125 ± 0.158 | 583.21 |
| UNet PhenDiff | 0.189 ± 0.025 | 09.64 ± 0.64 | 0.613 ± 0.531 | 62.50 |
| PertImage | 0.187 ± 0.095 | 11.46 ± 0.61 | 0.509 ± 0.043 | 46.88 |
| PertFlow | **0.205 ± 0.097** | **11.66 ± 0.68** | **0.511 ± 0.038** | **24.06** |

PhenDiff (12) is a conditional diffusion model that performs image-to-image translation to identify phenotypic shifts in microscopy images. The model operates through two stages: an inversion phase that maps real source images to Gaussian latent representations using DDIM deterministic sampling, followed by a generation phase that synthesizes images in the target condition. The UNet-based PhenDiff baseline predicts treatment-induced morphological changes without incorporating transcriptional information, serving as a purely vision-based approach. When evaluated on the GDPx3 dataset, UNet PhenDiff achieved moderate performance with SSIM of 0.189±0.025, PSNR of 9.64±0.64, and FID of 62.50, substantially outperforming the vanilla UNet baseline (SSIM 0.085±0.013, FID 583.21) but lagging behind multi-modal approaches. Deterministic UNet baseline performed even worse since it is trained with a simple MSE loss.

Table 3 shows equal loss weighting (0.5–0.5) achieves the best trade-off. Higher *w*_rna_ improves RNA accuracy but worsens image quality (higher FID). Higher *w*_img_ improves SSIM/PSNR but reduces RNA correlation metrics. Effect of KG weight *α*_drug_ = 0.3 is optimal. Smaller values (0.1, 0.2) underuse the biological prior, while larger values (0.4, 0.5) introduce noise that harms both modalities. Figure 4 shows similarity rating by ACVP board certified pathologists in blind review (10-point scale of similarity regarding morphology, detail and plausibility, with respect to ground truth Cell Painting images under corresponding chemical perturbations; 0 indicates poorest and 10 indicates the best). Two pathologists reported median score of 7.11 and 7.89 for PertFlow generated images, outperforming the baseline method PhenDiff (12) (median scores of 6.11 and 6.33), which confirmed the overall satisfying quality of cellular morphology images generated by PertFlow. Step-wise comparison (Figure 4) highlights the difference in inference dynamics. PertFlow generates recognizable structures by NFE 10 showing nuclear boundaries and cytoplasmic organization with near-final morphology. PertDiff requires more steps to reach similar organization. Both methods preserve multi-channel fluorescence distributions.

**Table 3.** Ablation study on loss weight ratios and knowledge graph embedding strength.

| $w_{\text{rna}}$ | $w_{\text{img}}$ | MSE ↓ | Pearson $r \uparrow$ | Spearman $r \uparrow$ | SSIM ↑ | PSNR ↑ | FID ↓ |
| --- | --- | --- | --- | --- | --- | --- | --- |
| 0.4 | 0.6 | 0.268 ± 0.752 | 0.761 ± 0.175 | 0.773 ± 0.047 | **0.215 ± 0.094** | **11.94 ± 0.65** | **21.73** |
| 0.5 | 0.5 | 0.231 ± 0.708 | 0.780 ± 0.264 | 0.792 ± 0.041 | 0.205 ± 0.097 | 11.66 ± 0.68 | 24.06 |
| 0.6 | 0.4 | **0.219 ± 0.694** | **0.791 ± 0.251** | **0.801 ± 0.038** | 0.193 ± 0.101 | 11.29 ± 0.73 | 27.15 |
| $\alpha_{\text{drug}}$ | | | | | | | |
| 0.1 | | 0.289 ± 0.784 | 0.741 ± 0.287 | 0.753 ± 0.052 | 0.187 ± 0.103 | 11.12 ± 0.76 | 29.31 |
| **0.3** |  | **0.231 ± 0.708** | **0.780 ± 0.264** | **0.792 ± 0.041** | **0.205 ± 0.097** | **11.66 ± 0.68** | **24.06** |
| 0.5 | | 0.248 ± 0.761 | 0.759 ± 0.279 | 0.769 ± 0.048 | 0.194 ± 0.101 | 11.38 ± 0.73 | 28.47 |

**Fig. 4.**
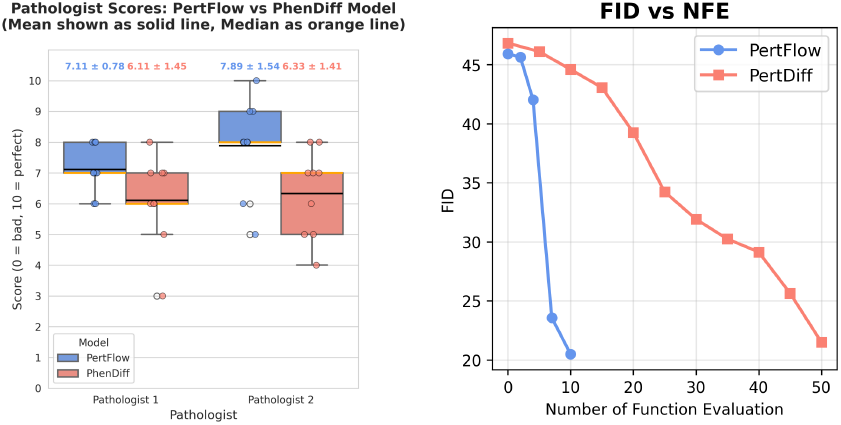
(LEFT) Pathologist image similarity score of generated vs. ground truth cellular morphology after treatment. Mean shown as solid line, median as dashed line. (RIGHT) FID versus number of function evaluations (NFE) across diffusion variants.

### Recovering drug-induced phenotype and morphology

Figure 1 shows the comparison of generated treatment vs real treatment images with drug name and concentration. Yellow boxes indicate similar cellular features due to drug perturbations in real and generated images. From left to right the drugs have the following effect on the cellular morphology: (1) Cevipabulin is a microtubule-destabilizing agent that binds to tubulin (26), disrupting microtubule dynamics, which leads to mitotic arrest and apoptosis in cancer cells. It shows anti-proliferative effects by inhibiting microtubule polymerization. (2) S-Camptothecin and its stereoisomers inhibit DNA topoisomerase (27), causing DNA damage during replication. This leads to DNA double-strand breaks, S-phase cell cycle arrest, and apoptosis, especially in rapidly dividing cells. (3) Podophyllotoxin binds to tubulin and inhibits microtubule assembly (28), resulting in mitotic arrest at metaphase and subsequent apoptosis. It serves as a precursor for etoposide, a topoisomerase II inhibitor. (4) Dabrafenib selectively inhibits mutant BRAF kinase (commonly V600E mutation) (29), blocking MAPK/ERK signaling pathway, leading to decreased tumor cell proliferation and inducing apoptosis in BRAF-mutated cancer cells.

ACVP certified pathologists provided descriptions of generated and real treatment image results: In Figure 5 control images exhibited typical fusiform cells with parallel alignment and organized architecture. Both predicted and real Ouabain (0.3 *µ*M) treatment showed decreased cellular density, reduced cell size, increased intercellular spacing, nuclear condensation, disrupted parallel orientation with cells. Similarly, predicted and actual Podophyllotoxin (3.0 *µ*M) treatment displayed increased intercellular spacing, elevated multi-nucleated cell populations indicative of cellular injury, loss of fusiform morphology, and decreased cellular density. Suprafenacine (3.0 *µ*M) predictions and actual treatment both revealed cellular disorganization, loss of fusiform shape, cellular fragmentation, nuclear size reduction, decreased cell-to-cell contact.

**Fig. 5.**
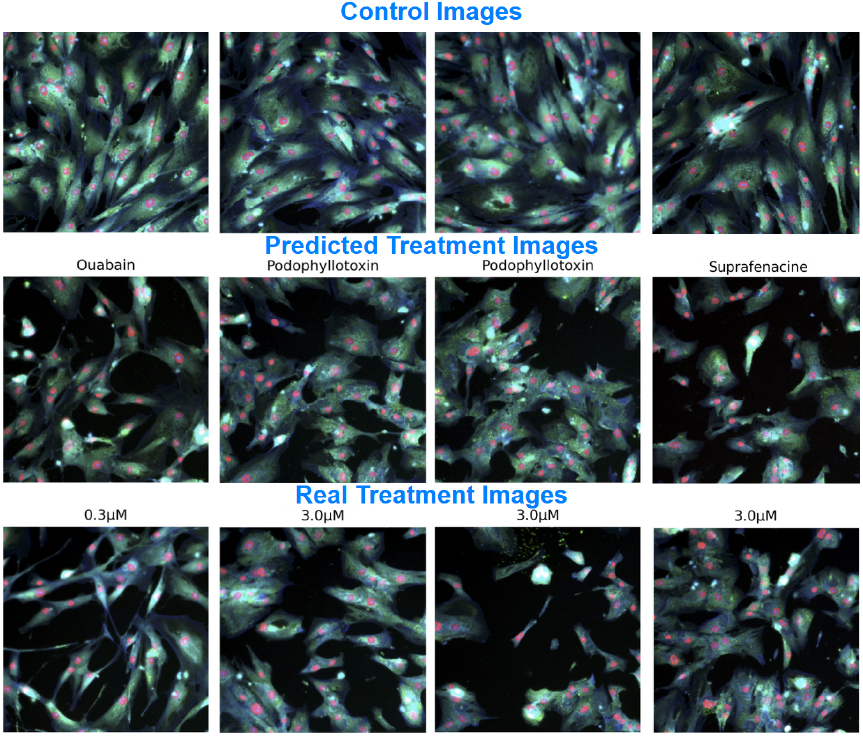
Generated vs real treatment for Aortic Smooth Muscle Cell

In Figure 6 control images displayed characteristic fibroblast morphology with fusiform cells, organized cellular arrangement, and appropriate cell-to-cell contact. Both predicted and real Podophyllotoxin (0.3 *µ*M) treatment exhibited decreased cellular density, reduced cell-to-cell contact, and loss of fusiform morphology while maintaining nuclear size. Panobinostat (3.0 *µ*M) predictions and actual treatment showed minimal morphological deviation from control despite the higher dosage, with the model correctly preserving the relatively unaltered cellular architecture. Nocodazole treatment at 3.0 *µ*M demonstrated strong concordance between predicted and real images, both displaying substantial loss of cellular density and fusiform morphology, with cells adopting rounded, blob-like shapes while nuclear size remained similar. At the lower nocodazole dose (0.3 *µ*M), both predicted and actual treatments showed attenuated phenotypic changes including decreased cellular density, partial retention of fusiform morphology, reduced cellular elongation, less pronounced blob-like transformation compared to the higher dose.

**Fig. 6.**
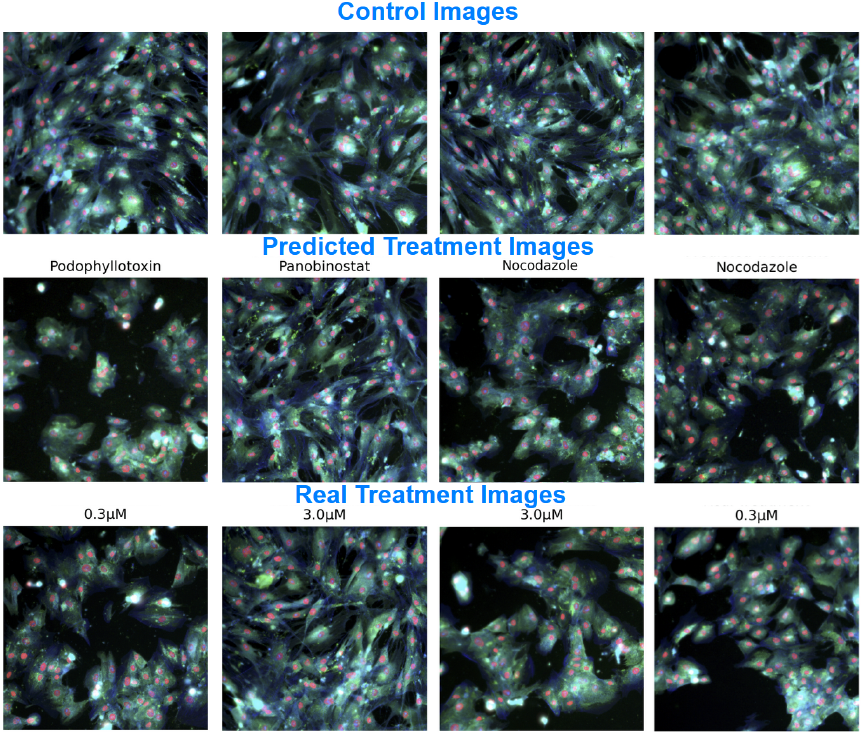
Generated vs real treatment for Dermal Fibroblasts.

In Figure 7 control images exhibited typical A549 morphology with appropriate cellular density and organization. DMSO treatment (0.0 *µ*M) served as vehicle control, with both predicted and real images showing no morphological deviation from control conditions. Nocodazole (1.0 *µ*M) predictions and actual treatment both displayed preserved cellular morphology with substantially decreased cellular density and reduced cellularity. Emetine dihydrochloride hydrate (1.0 *µ*M) showed concordance between predicted and real images, both exhibiting decreased cellular density and diminished cellular cohesiveness. Torin2 (0.1 *µ*M) predictions and actual treatment demonstrated increased angular cytoplasmic projections, moderate reduction in cellular density, and decreased cellular cohesion. The morphological agreement between predicted and real treatment conditions across varying drug concentrations validates the model’s dose-dependent prediction capability. Further results of generated images for all drug compounds can be found in the appendix (Figures 12, 13, 14, 15, 16).

**Fig. 7.**
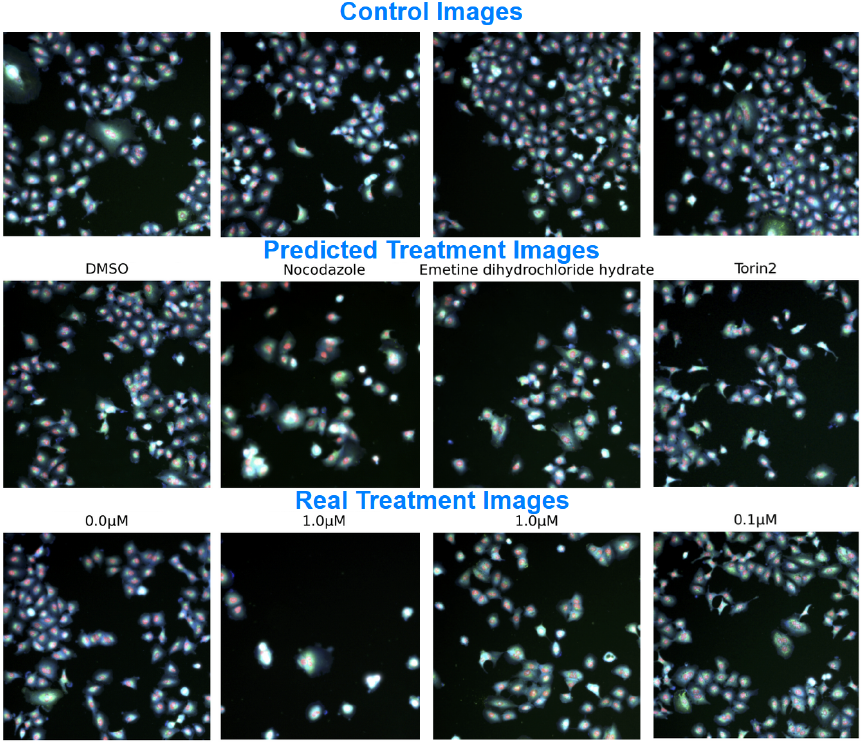
Generated vs real treatment for A549.

Figure 2 UMAPs (30) demonstrate PertFlow’s biological coherence in generating treatment responses. Left panel shows clear control-treatment cluster separation, indicating distinct transcriptomic signatures captured in embedding space. Right panel shows generated treatment samples clustering with real samples, particularly in high-response regions, demonstrating that cross-modal attention effectively leverages imaging to predict directional gene expression changes and produce biologically realistic treatment profiles.

Figure 8, 9, 10 UMAP and t-SNE illustrate cross-modal attention’s effect on feature alignment. The top panels show transformation from separate feature spaces (left) to joint aligned space (middle), with RNA-seq (blue) and image features (red) demonstrating successful correspondence learning. The middle and bottom panels reveal that cross-modal attention transforms scattered, unstructured distributions into highly structured, clustered organizations in both modalities, indicating that attention not only aligns features across modalities but enhances the internal structure and discriminative power of each individual feature space. The shared representation space (right) demonstrates successful integration of RNA, imaging, and drug modalities. Compound-based organization indicates that cross-modal attention effectively combines gene expression, imaging, and drug conditioning into a unified latent space where biologically similar samples cluster together regardless of modality, enabling improved treatment response prediction and generation.

**Fig. 8.**
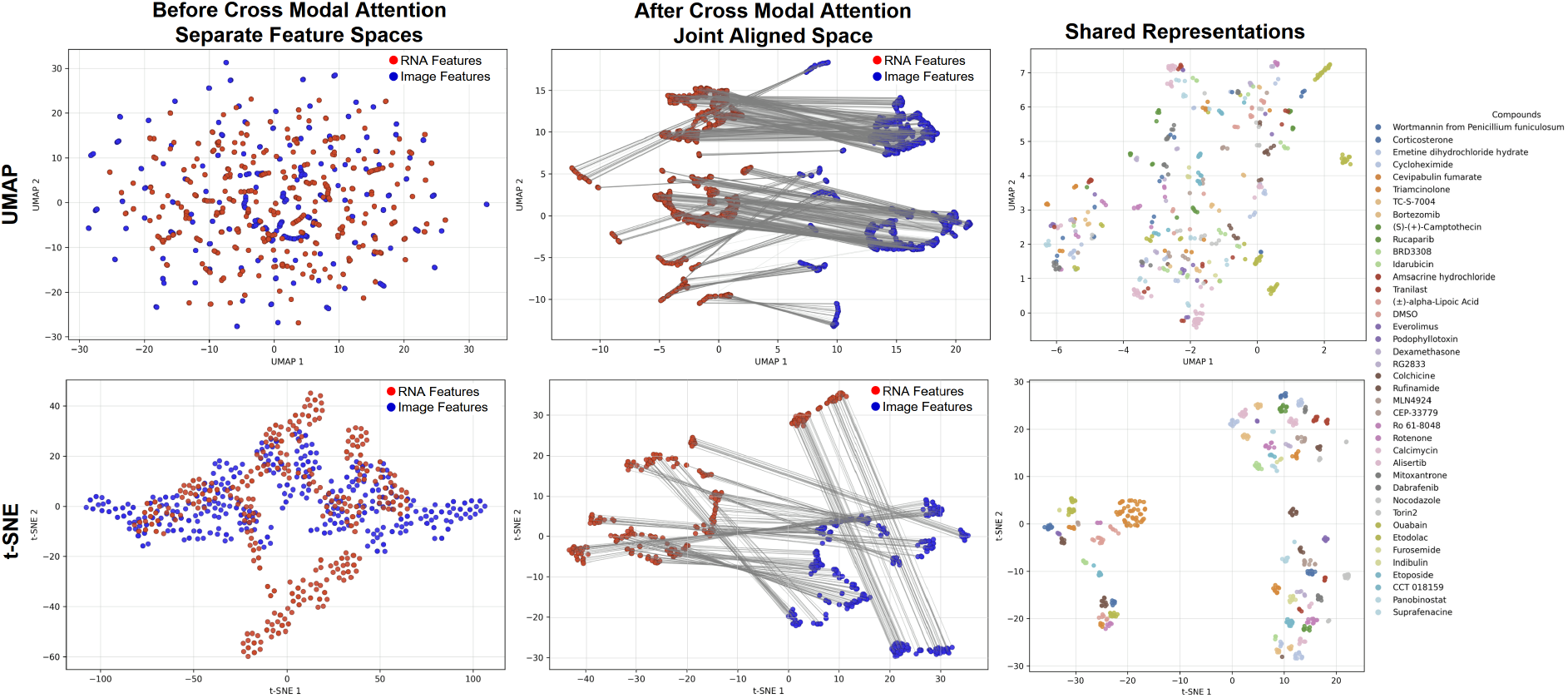
UMAP and t-SNE of RNA and image embeddings before and after cross modal attention

**Fig. 9.**
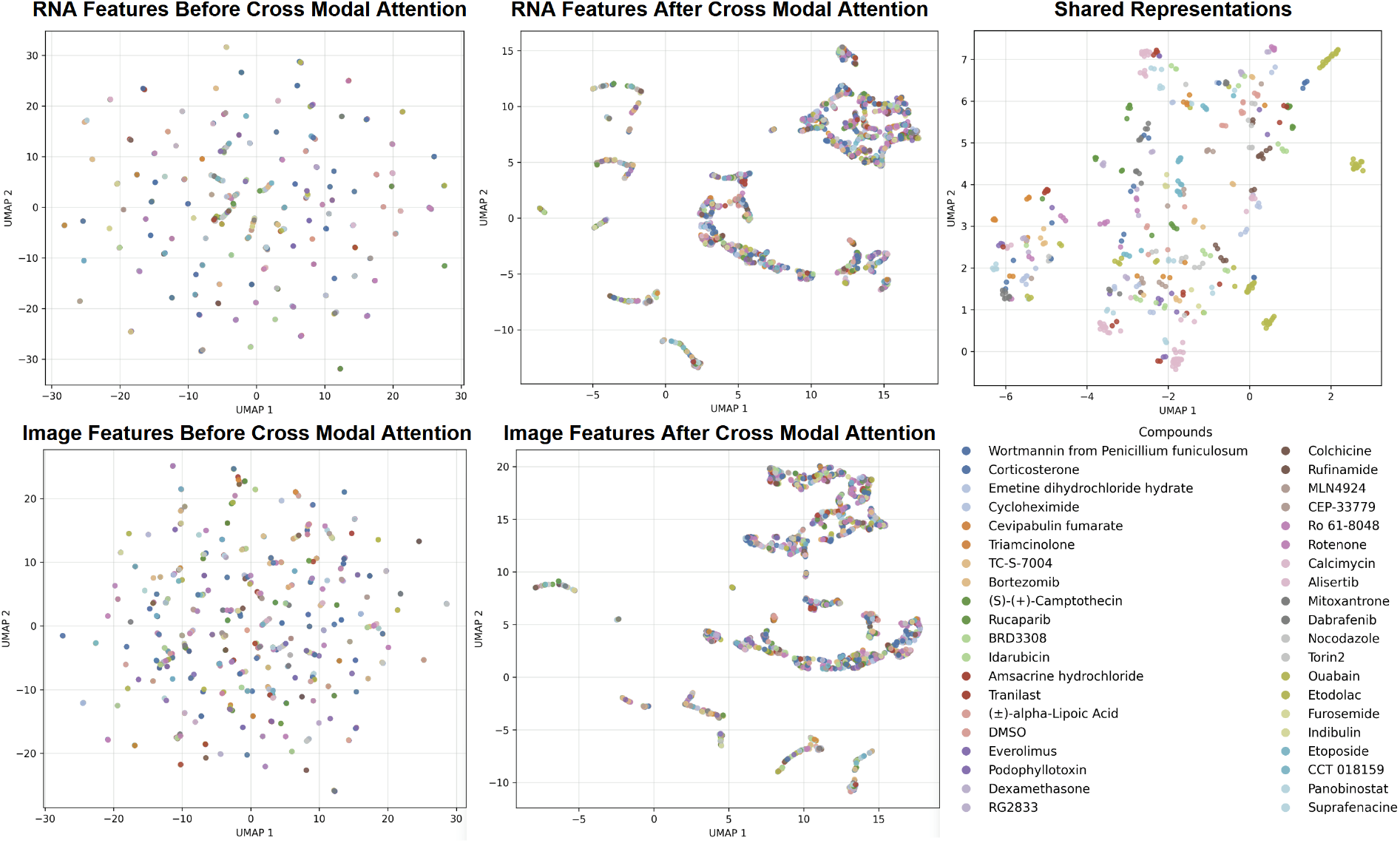
UMAP of RNA and image embeddings before and after cross modal attention block

**Fig. 10.**
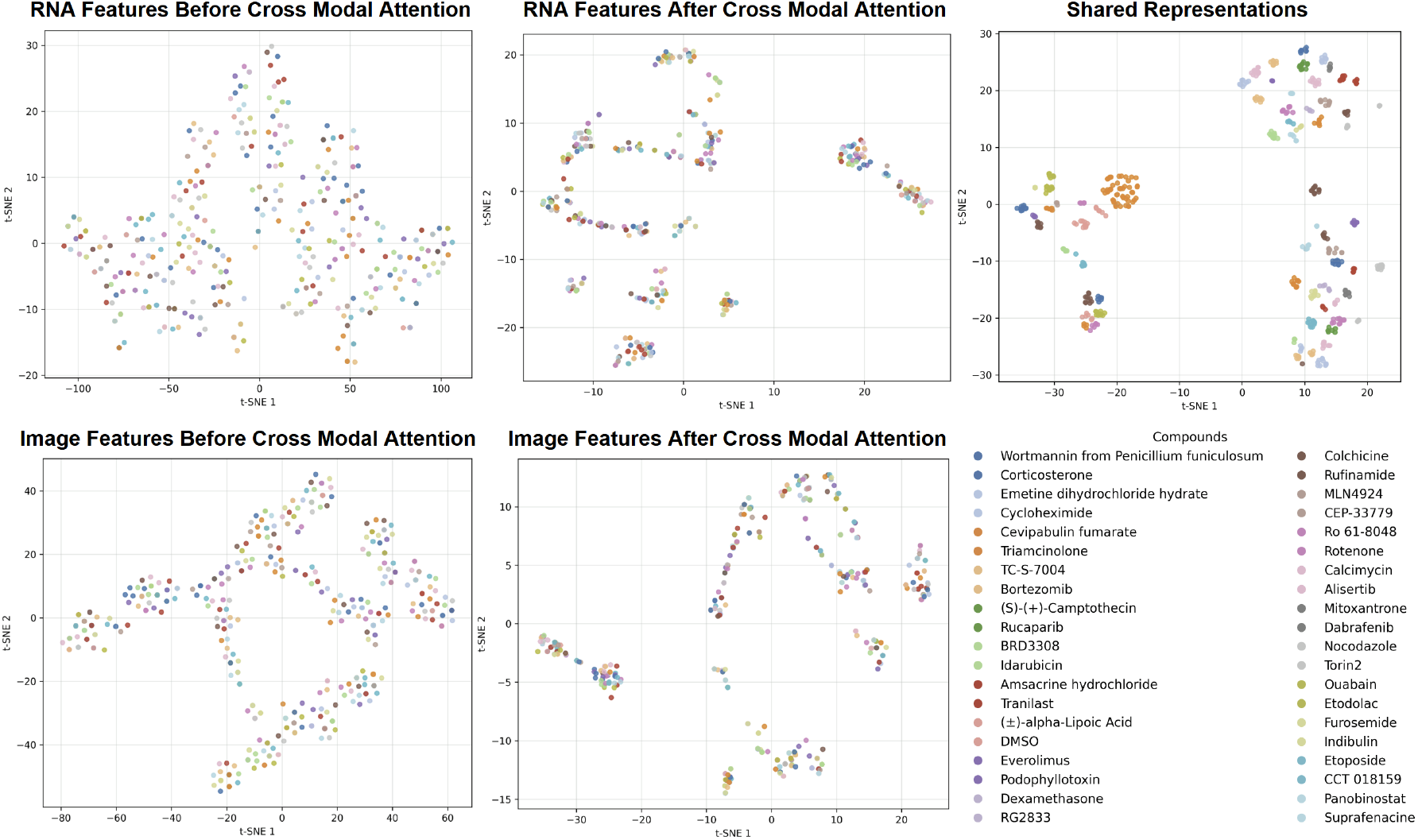
t-SNE of RNA and image embeddings before and after cross modal attention block

Cross-modal attention shows correspondence between RNA-image modalities, preserving distinct structures. Pairwise alignment distances average 16.48 (UMAP) and 37.44 (t-SNE), with positive modality separation scores (UMAP: 0.81, t-SNE: 0.59) confirming preserved modality-specific features. Compound clustering improves substantially, with silhouette scores rising from -0.74 to -0.20 (UMAP) and -0.81 to -0.28 (t-SNE). The shared space separation ratio of 1.33 validates treatment-specific representations consistent across both modalities. Alignment metrics used standardized embeddings projected into 2D via UMAP and t-SNE. Pairwise alignment distances were averaged Euclidean distances between corresponding RNA-image pairs. Modality separation used silhouette scores with binary labels (0=RNA, 1=Image), where positive values indicate preserved modality-specific structures. Compound clustering quality was assessed via silhouette scores with treatment labels. Shared space separation ratio was calculated as mean inter-compound cosine distance divided by mean intra-compound distance, values *>*1 indicate successful treatment discrimination.

### Gene Enrichment Analysis

To evaluate whether PertFlow learns mechanistically accurate representations without explicit pathway supervision, we extracted the input RNA gradient magnitudes (|∇ _*x*_*L*|) with respect to the generated image velocity shown in Figure 11. Gene Set Enrichment Analysis (GSEA) on these gradients across a diversity panel of 10 canonical drugs reveals that the model accurately prioritizes genes mapping directly to known mechanisms of action. For instance, the microtubule destabilizer Colchicine strongly drives MAPK stress signaling, while the corticosteroid Corticosterone leverages NF-*κ*B signaling. Dot size represents statistical significance (−log_10_(FDR)) and color indicates the Normalized Enrichment Score (NES). Generic cancer umbrellas were filtered to highlight specific mechanism-of-action pathways.

**Fig. 11.**
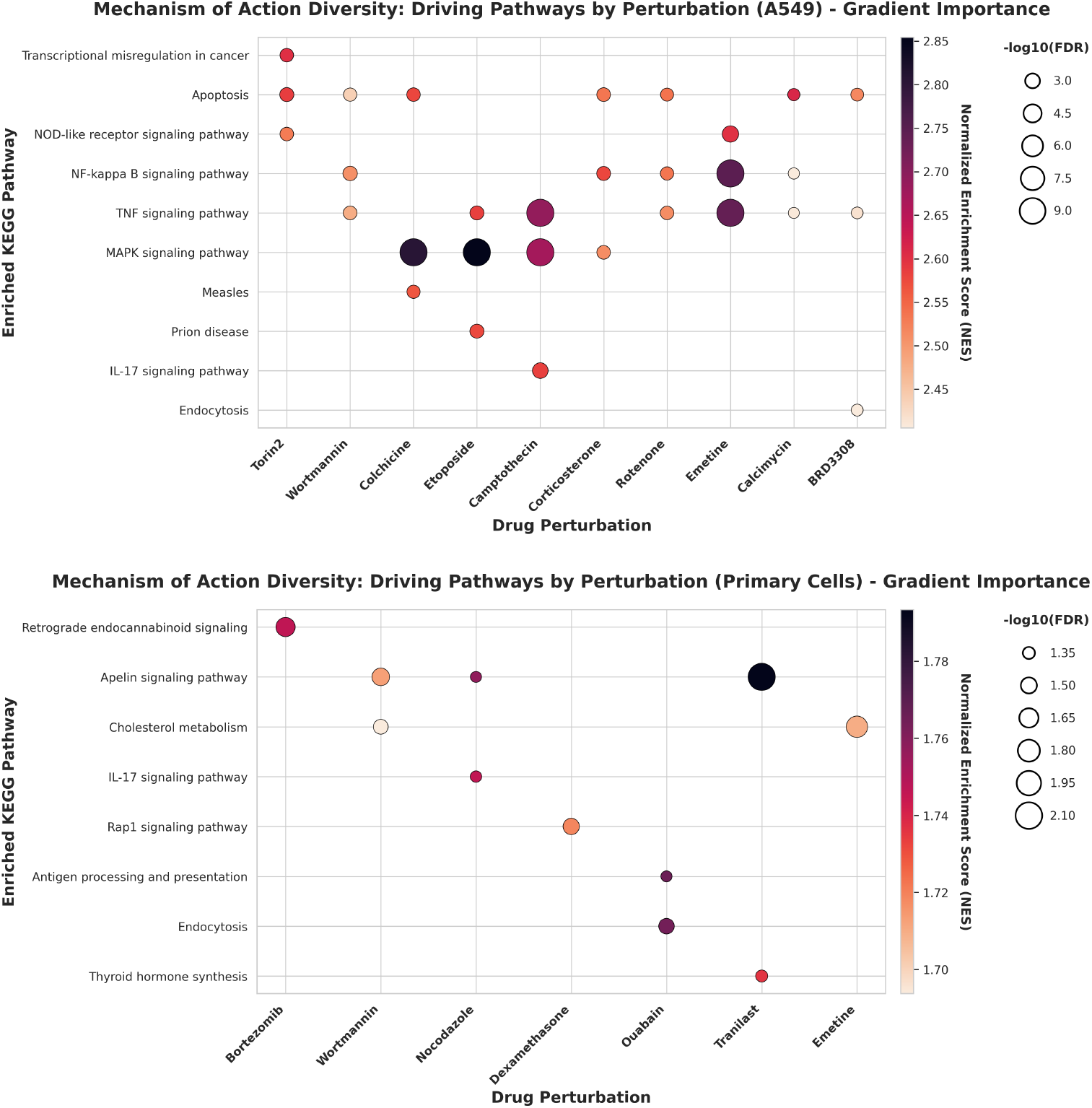
Zero-shot discovery of perturbation mechanisms via generative gradients

## Discussion

PertFlow represents a foundational step toward unified modeling of multi-modal cellular drug responses, bridging the molecular (transcriptomic) and phenotypic (morphological) effects of chemical perturbations. Unlike previous approaches that treat these modalities in isolation or only predict one from the other, PertFlow achieves simultaneous, drug-conditioned generation of both gene expression profiles and cellular morphology. The integration of control transcriptomic and imaging data into a shared embedding space, combined with rectified flow dynamics, enables bio-logically consistent synthesis of treatment outcomes. Despite this progress, several challenges remain. First, generalization to unseen cell lines or novel compounds is limited by the scarcity of paired multi-modal datasets with shared metadata. While PertFlow can still infer morphological changes from control data alone, future work should explore more targeted integration of chemogenomic databases (e.g., ChEMBL, DrugBank) that explicitly encode compound-target binding affinities and structure-activity relationships, building upon our current PrimeKG integration which provides broader biological context but lacks fine-grained chemical similarity information critical for generalizing to truly novel compounds. Second, while PertFlow enables modality translation, aligning embeddings across modalities may inadvertently entangle task-relevant factors. Disentangling causal latent factors remains an open question for cross-modal modeling. Third, the in vitro context of our experiments may not capture drug effects requiring complex microenvironmental interactions, such as immune modulation. Extending PertFlow to model cell-cell communication or tissue-level organization could enhance translational utility. Overall, PertFlow sets the stage for future cross-modal generative modeling in drug discovery, offering a unified framework for understanding how molecular mechanisms manifest as observable phenotypes under pharmacological perturbation.

## Conclusion

We introduced PertFlow, a unified generative framework for jointly modeling transcriptomic and morphological drug responses using cross-modal attention and rectified flow dynamics. By aligning control RNA-seq and image features through a shared embedding space conditioned on drug meta-data, PertFlow enables simultaneous prediction of treatment gene expression and synthesis of cellular morphology. Extensive evaluation on the GDPx3 dataset demonstrates strong cross-modal consistency, biologically realistic image generation, and competitive transcriptomic prediction performance, outperforming single-modality and diffusion baselines. UMAP analysis of cross-modal attention and shared embeddings further strengthen our hypothesis. Our results highlight PertFlow’s potential for virtual drug screening, mechanistic hypothesis generation, and multi-modal perturbation analysis, paving the way for more integrative and interpretable approaches to pharmacological modeling.

## Supplementary Note A: Appendix

### A. Architecture Details

Our RNA-seq encoder processes gene expression data through multi-layer self-attention (31) to capture gene-gene interactions:

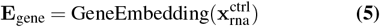

where each gene expression value is projected to a *d*_gene_-dimensional embedding space. We apply *L* layers of multi-head self-attention (MHA):

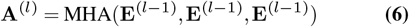

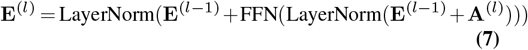

The final RNA-seq features are obtained through attention-based pooling:

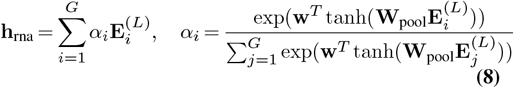

Control cellular images are processed through a ResNet-style convolutional (32) architecture:

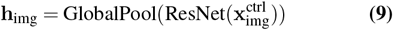

Drug conditioning information combines categorical and continuous variables:

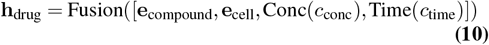

where **e**_compound_ and **e**_cell_ are learned embeddings for compound and cell line identities.

Knowledge graph integration enhances both molecular and genomic representations through structured biological knowledge from PrimeKG (33). For drug embeddings, compounds are mapped to knowledge graph entities capturing molecular interactions, pathways, and pharmacological relationships. The heterogeneous graph neural network processes drug-protein, drug-drug, and protein-protein interactions:

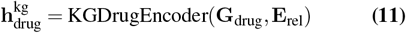

where **G**_drug_ represents drug nodes and **E**_rel_ captures multi-relational edges. Similarly, gene expressions are enhanced with protein interaction networks and pathway information:

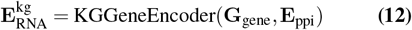

The knowledge graph embeddings are integrated additively with learned representations:

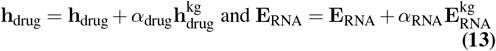

where *α*_drug_ = 0.3 and *α*_RNA_ = 0.3 are learned weighting factors.

To capture cross-modal RNA-Image dependencies, we use multi-token cross-attention. Each modality is projected to *K* token representations:

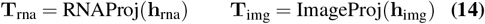

Each modality goes through a self-attention block, then cross-attention is applied bidirectionally (Eq. 6):

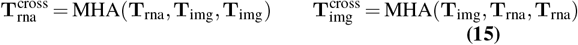

Enhanced features are obtained through residual connections and attention pooling:

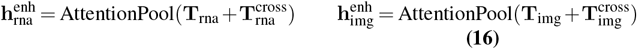

The cross-modal features are combined with drug conditioning (Eq. 10) to form a unified representation:

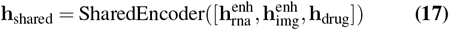

This shared representation captures the complex dependencies between molecular states, morphological features, and drug effects necessary for coherent multi-modal generation. Treatment gene expression is generated through direct prediction from the shared representation (Eq. 17):

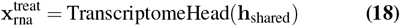

For image generation, we adapt rectified flow dynamics. Given noise **z**_0_ ∼ 𝒩 (0, **I**) and target image 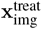, rectified flow defines a linear interpolation path:

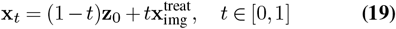

The velocity field is defined as:

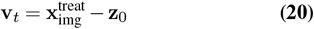

and our multi-modal-conditioned UNet learns to predict this velocity:

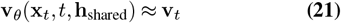

The UNet incorporates cross-attention layers that attend to image conditioning derived from the shared representation:

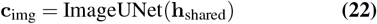

Our training strategy combines task-specific losses with cross-modal consistency objectives. We use a combination of MSE and auxiliary Pearson Correlation loss for transcriptome prediction:

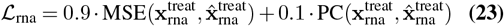

For rectified flow training, we minimize the velocity prediction error for image generation:

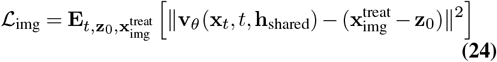

We implement triplet contrastive consistency (34) to ensure well-aligned features produce better predictions than mis-aligned ones:

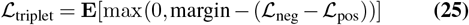

where ℒ_pos_ is the prediction error with aligned features and ℒ_neg_ with misaligned features. The complete training objective combines all losses (Eqs. 23-25):

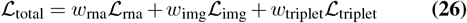

where the weights are set to *w*_rna_ = 0.5, *w*_img_ = 0.5, *w*_triplet_ = 0.05.

Treatment RNA-seq is generated through a single forward pass using Eq. 18:

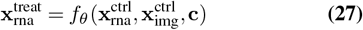

For high-quality image generation, we use an adaptive DO-PRI5 solver (35) that iteratively integrates the learned velocity field:

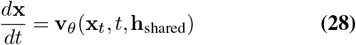

Starting from noise **x**_0_ ∼ 𝒩(0, **I**) at *t* = 0, the solver adaptively adjusts step sizes based on error estimation to reach the treatment image at *t* = 1. The adaptive integration ensures both computational efficiency and generation quality. The DOPRI5 method uses a 5th-order Runge-Kutta scheme with embedded 4th-order error estimation for automatic step size control. The step size *h* is adapted based on the estimated local truncation error to maintain tolerance levels.

### B. Further Examples of Generated Images

**Fig. 12.**
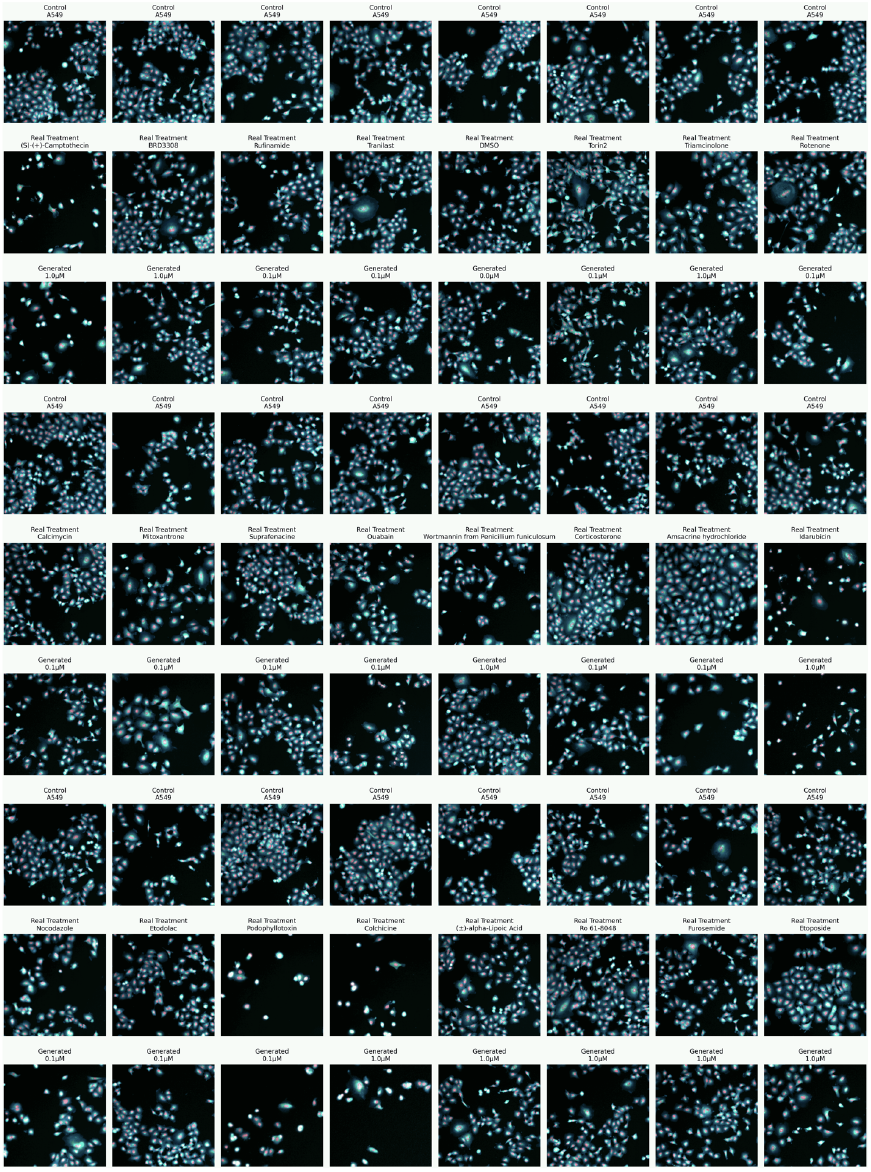
Cell line A549

**Fig. 13.**
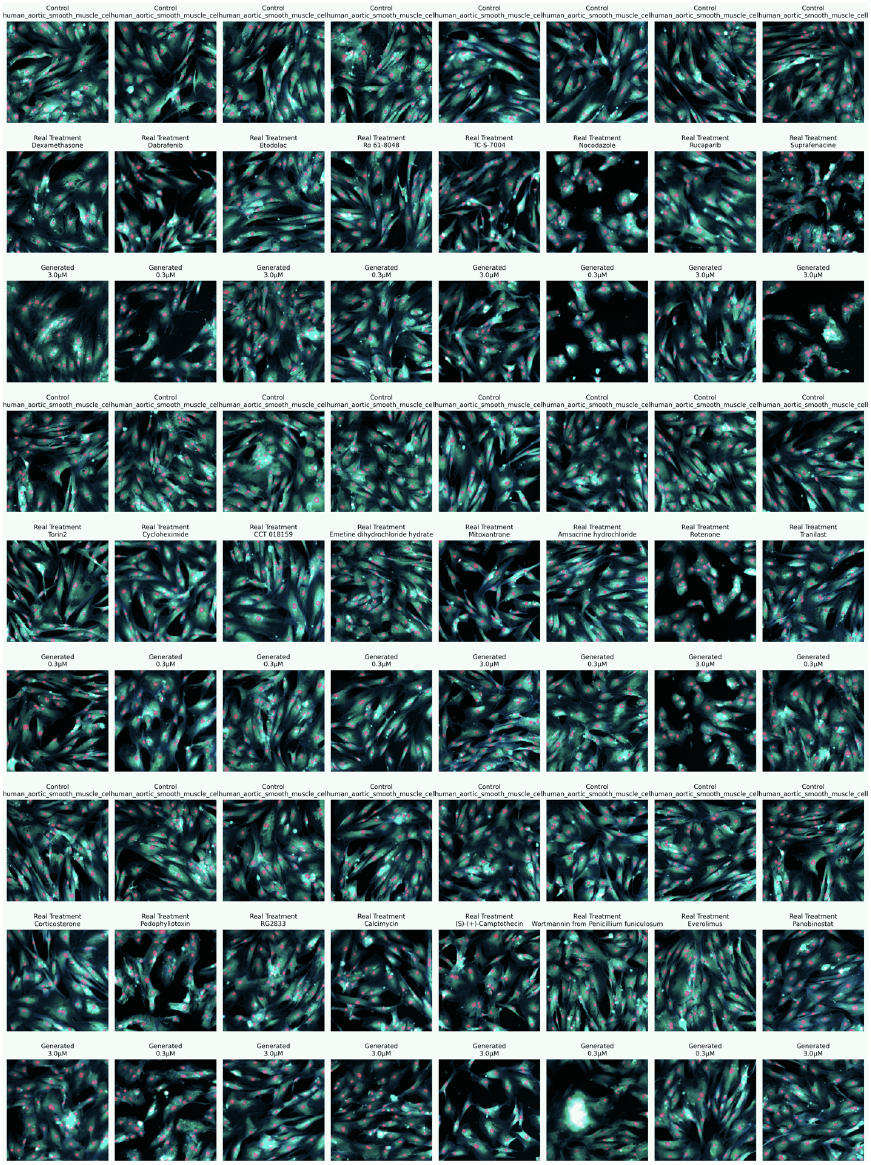
Cell line Aortic Smooth Muscle Cell

**Fig. 14.**
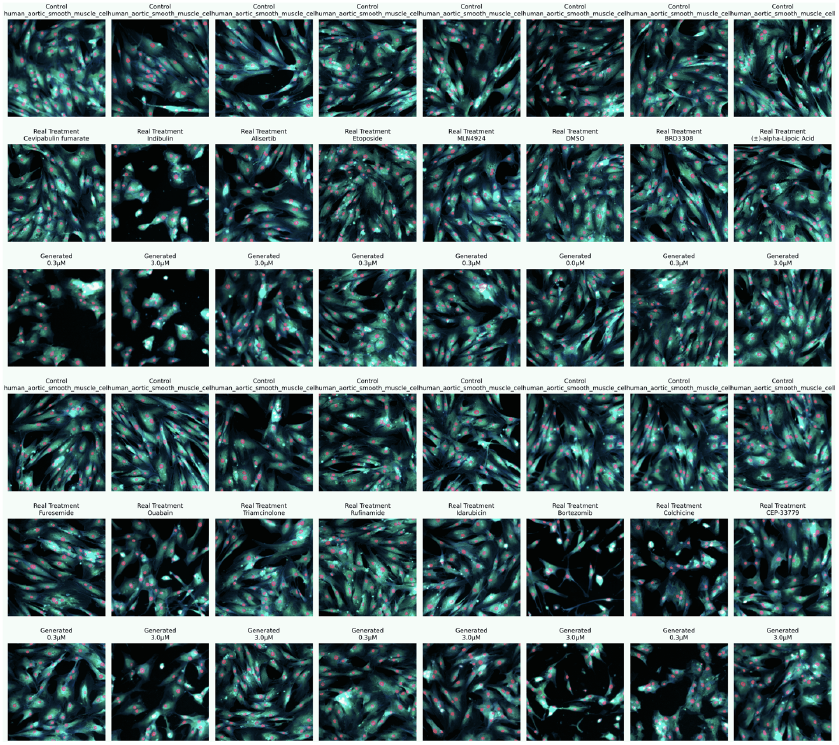
Cell line Aortic Smooth Muscle Cell

**Fig. 15.**
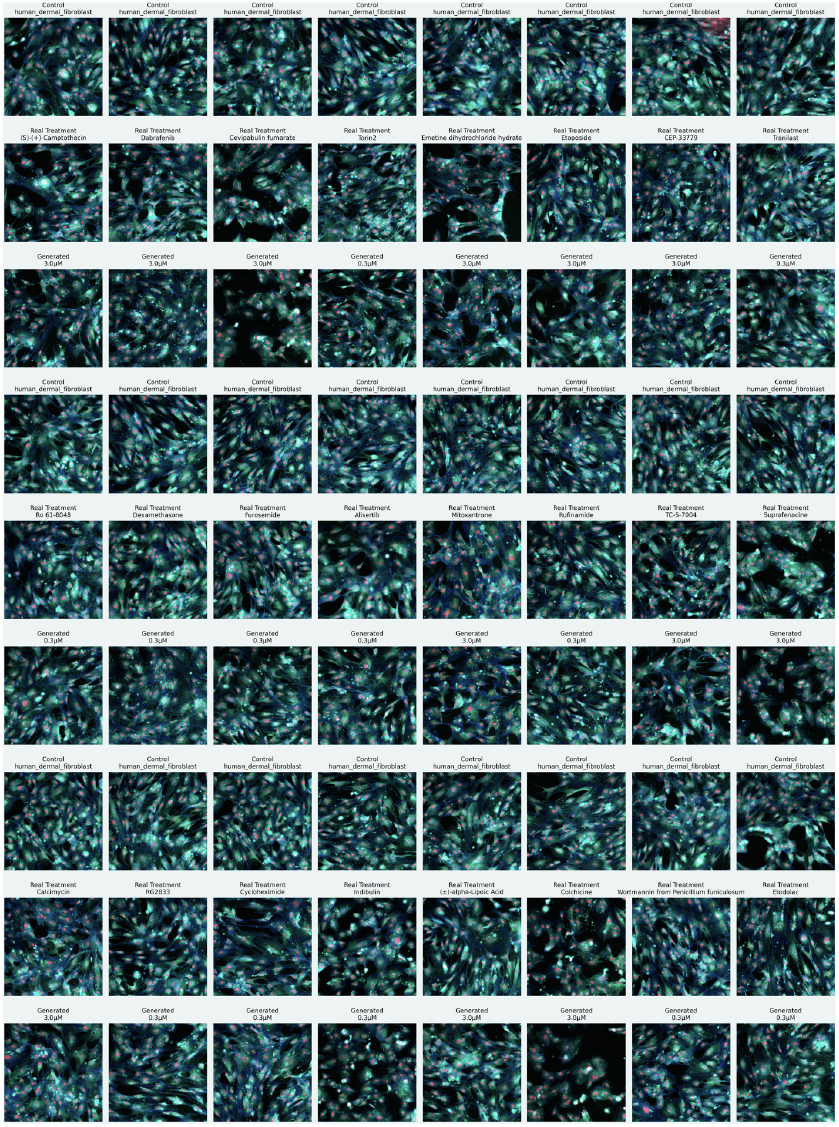
Cell line Dermal Fibroblast

**Fig. 16.**
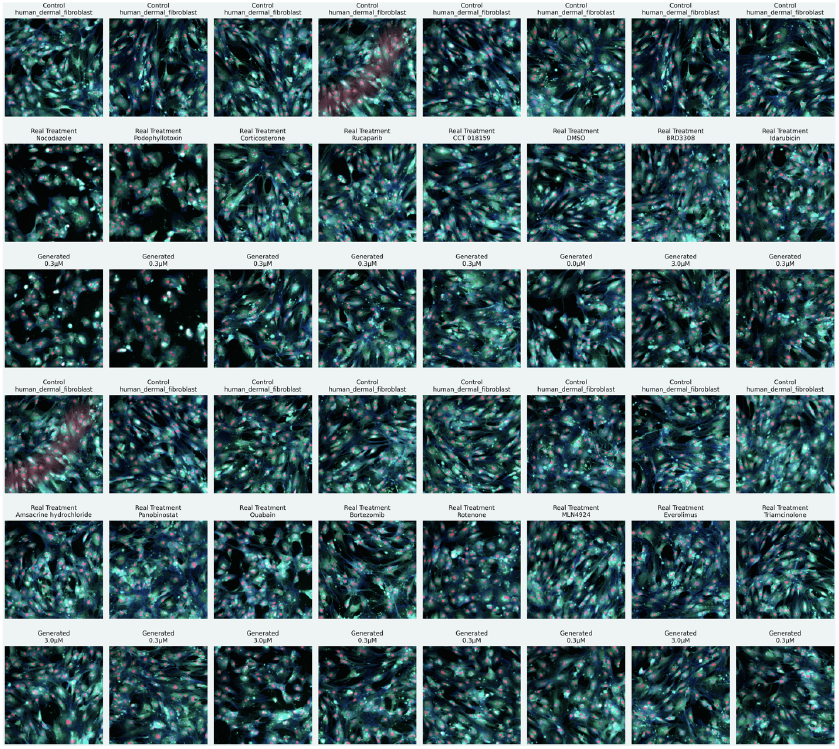
Cell line Dermal Fibroblast

## References

1. Mohammad Lotfollahi, F Alexander Wolf, and Fabian J Theis. scgen predicts single-cell perturbation responses. Nature methods, 16(8):715–721, 2019.

2. Mohammad Lotfollahi, Anna Klimovskaia Susmelj, Carlo De Donno, Yuge Ji, Ignacio L Ibarra, F Alexander Wolf, Nafissa Yakubova, Fabian J Theis, and David Lopez-Paz. Compositional perturbation autoencoder for single-cell response modeling. BioRxiv, 2021.

3. Leon Hetzel, Simon Boehm, Niki Kilbertus, Stephan Günnemann, Fabian Theis, et al. Predicting cellular responses to novel drug perturbations at a single-cell resolution. Advances in Neural Information Processing Systems, 35:26711–26722, 2022.

4. Yusuf Roohani, Kexin Huang, and Jure Leskovec. Predicting transcriptional outcomes of novel multigene perturbations with gears. Nature Biotechnology, 42(6):927–935, 2024.

5. Xiaoning Qi, Lianhe Zhao, Chenyu Tian, Yueyue Li, Zhen-Lin Chen, Peipei Huo, Runsheng Chen, Xiaodong Liu, Baoping Wan, Shengyong Yang, et al. Predicting transcriptional responses to novel chemical perturbations using deep generative model for drug discovery. Nature Communications, 15(1):9256, 2024.

6. Xiaochu Tong, Ning Qu, Xiangtai Kong, Shengkun Ni, Jingyi Zhou, Kun Wang, Lehan Zhang, Yiming Wen, Jiangshan Shi, Sulin Zhang, et al. Deep representation learning of chemical-induced transcriptional profile for phenotype-based drug discovery. Nature Communications, 15(1):5378, 2024.

7. Constantin Ahlmann-Eltze, Wolfgang Huber, and Simon Anders. Deep-learning-based gene perturbation effect prediction does not yet outperform simple linear baselines. Nature Methods, pages 1–5, 2025.

8. Mark-Anthony Bray, Shantanu Singh, Han Han, Chadwick T Davis, Blake Borgeson, Cathy Hartland, Maria Kost-Alimova, Sigrun M Gustafsdottir, Christopher C Gibson, and Anne E Carpenter. Cell painting, a high-content image-based assay for morphological profiling using multiplexed fluorescent dyes. Nature protocols, 11(9):1757–1774, 2016.

9. Claire McQuin, Allen Goodman, Vasiliy Chernyshev, Lee Kamentsky, Beth A Cimini, Kyle W Karhohs, Minh Doan, Liya Ding, Susanne M Rafelski, Derek Thirstrup, et al. Cellprofiler 3.0: Next-generation image processing for biology. PLoS biology, 16(7):e2005970, 2018.

10. Carsen Stringer, Tim Wang, Michalis Michaelos, and Marius Pachitariu. Cellpose: a generalist algorithm for cellular segmentation. Nature methods, 18(1):100–106, 2021.

11. Qiaosi Tang, Ranjala Ratnayake, Gustavo Seabra, Zhe Jiang, Ruogu Fang, Lina Cui, Yousong Ding, Tamer Kahveci, Jiang Bian, Chenglong Li, et al. Morphological profiling for drug discovery in the era of deep learning. Briefings in Bioinformatics, 25(4), 2024.

12. Anis Bourou, Thomas Boyer, Kévin Daupin, Véronique Dubreuil, Aurélie De Thonel, Valérie Mezger, and Auguste Genovesio. Phendiff: Revealing invisible phenotypes with conditional diffusion models. CoRR, 2023.

13. Alexis Lamiable, Tiphaine Champetier, Francesco Leonardi, Ethan Cohen, Peter Sommer, David Hardy, Nicolas Argy, Achille Massougbodji, Elaine Del Nery, Gilles Cottrell, et al. Revealing invisible cell phenotypes with conditional generative modeling. Nature Communications, 14(1):6386, 2023.

14. Ronald Xie, Kuan Pang, Sai Chung, Catia Perciani, Sonya MacParland, Bo Wang, and Gary Bader. Spatially resolved gene expression prediction from histology images via bi-modal contrastive learning. Advances in Neural Information Processing Systems, 36:70626– 70637, 2023.

15. Charles Comiter. Inference of single cell profiles from histology stains with the Single-Cell omics from Histology Analysis Framework (SCHAF). Massachusetts Institute of Technology, 2024.

16. Chongyue Zhao, Zhongli Xu, Xinjun Wang, Shiyue Tao, William A MacDonald, Kun He, Amanda C Poholek, Kong Chen, Heng Huang, and Wei Chen. Innovative super-resolution in spatial transcriptomics: a transformer model exploiting histology images and spatial gene expression. Briefings in Bioinformatics, 25(2):bbae052, 2024.

17. Reuben A Saunders, William E Allen, Xingjie Pan, Jaspreet Sandhu, Jiaqi Lu, Thomas K Lau, Karina Smolyar, Zuri A Sullivan, Catherine Dulac, Jonathan S Weissman, et al. Perturb-multimodal: A platform for pooled genetic screens with imaging and sequencing in intact mammalian tissue. Cell, 2025.

18. Löïc Binan, Aiping Jiang, Serwah A Danquah, Vera Valakh, Brooke Simonton, Jon Bezney, Robert T Manguso, Kathleen B Yates, Ralda Nehme, Brian Cleary, et al. Simultaneous crispr screening and spatial transcriptomics reveal intracellular, intercellular, and functional transcriptional circuits. Cell, 188(8):2141–2158, 2025.

19. Xiaohua Lu, Liangxu Xie, Lei Xu, Rongzhi Mao, Xiaojun Xu, and Shan Chang. Multimodal fused deep learning for drug property prediction: Integrating chemical language and molecular graph. Computational and Structural Biotechnology Journal, 23:1666–1679, 2024.

20. Zhiye Guo, Jian Liu, Yanli Wang, Mengrui Chen, Duolin Wang, Dong Xu, and Jianlin Cheng. Diffusion models in bioinformatics and computational biology. Nature reviews bioengineering, 2(2):136–154, 2024.

21. Patrick Esser, Sumith Kulal, Andreas Blattmann, Rahim Entezari, Jonas Müller, Harry Saini, Yam Levi, Dominik Lorenz, Axel Sauer, Frederic Boesel, et al. Scaling rectified flow transformers for high-resolution image synthesis. In Forty-first international conference on machine learning, pages 531–540, 2024.

22. Xingchao Liu, Chengyue Gong, and Qiang Liu. Flow straight and fast: Learning to generate and transfer data with rectified flow. arXiv preprint arXiv:2209.03003, 2022.

23. Yaron Lipman, Ricky TQ Chen, Heli Ben-Hamu, Maximilian Nickel, and Matt Le. Flow matching for generative modeling. arXiv preprint arXiv:2210.02747, 2022.

24. API Model and Enzymes Biologics. Ginkgo datapoints: Data generation for ai model training. Ginkgo Bioworks, 2025.

25. Huimin Huang, Lanfen Lin, Ruofeng Tong, Hongjie Hu, Qiaowei Zhang, Yutaro Iwamoto, Xianhua Han, Yen-Wei Chen, and Jian Wu. Unet 3+: A full-scale connected unet for medical image segmentation. In ICASSP 2020-2020 IEEE international conference on acoustics, speech and signal processing (ICASSP), pages 1055–1059. Ieee, 2020.

26. Jianhong Yang, Yamei Yu, Yong Li, Wei Yan, Haoyu Ye, Lu Niu, Minghai Tang, Zhoufeng Wang, Zhuang Yang, Heying Pei, et al. Cevipabulin-tubulin complex reveals a novel agent binding site on α-tubulin with tubulin degradation effect. Science Advances, 7(21): eabg4168, 2021.

27. Corwin Hansch and Rajeshwar P Verma. 20-(s)-camptothecin analogues as dna topoisomerase i inhibitors: a qsar study. ChemMedChem: Chemistry Enabling Drug Discovery, 2 (12):1807–1813, 2007.

28. Stephanie Desbene and Sylviane Giorgi-Renault. Drugs that inhibit tubulin polymerization: the particular case of podophyllotoxin and analogues. Current Medicinal Chemistry-Anti-Cancer Agents, 2(1):71–90, 2002.

29. David Planchard, Benjamin Besse, Harry JM Groen, Sayed MS Hashemi, Julien Mazieres, Tae Min Kim, Elisabeth Quoix, Pierre-Jean Souquet, Fabrice Barlesi, Christina Baik, et al. Phase 2 study of dabrafenib plus trametinib in patients with braf v600e-mutant metastatic nsclc: updated 5-year survival rates and genomic analysis. Journal of Thoracic Oncology, 17(1):103–115, 2022.

30. Leland McInnes, John Healy, and James Melville. Umap: Uniform manifold approximation and projection for dimension reduction. arXiv preprint arXiv:1802.03426, 2018.

31. Ashish Vaswani, Noam Shazeer, Niki Parmar, Jakob Uszkoreit, Llion Jones, Aidan N Gomez, Lukasz Kaiser, and Illia Polosukhin. Attention is all you need. In Advances in neural information processing systems, pages 5998–6008, 2017.

32. Kaiming He, Xiangyu Zhang, Shaoqing Ren, and Jian Sun. Deep residual learning for image recognition. In Proceedings of the IEEE conference on computer vision and pattern recognition, pages 770–778, 2016.

33. Payal Chandak, Kexin Huang, and Marinka Zitnik. Building a knowledge graph to enable precision medicine. Scientific Data, 10(1):67, 2023.

34. George Stoica, Vivek Ramanujan, Xiang Fan, Ali Farhadi, Ranjay Krishna, and Judy Hoffman. Contrastive flow matching. arXiv preprint arXiv:2506.05350, 2025.

35. John R Dormand and Peter J Prince. Runge-kutta triples. Computers & Mathematics with Applications, 12(9):1007–1017, 1986.

